# Decolonizing Psocopteran Systematics: Holarctic Lineages Cannot Inform Diversity and Evolution in Tropics

**DOI:** 10.1101/2020.10.02.324277

**Authors:** Valentina Sarria-Rodríguez, Ranulfo Gonzalez-Obando, Nelson Rivera-Franco, Heiber Cardenas-Henao, Cristian Román-Palacios

## Abstract

Despite tropical psocids comprise ~60% of species diversity within the Psocidae (Insecta, Psocodea), previous studies on the Psocidae phylogeny have poorly sampled tropical species (<40% species in trees). Here we discuss the evolution and systematics of the Psocidae based on the most comprehensive species-level sampling of the Psocidae. We sequenced and inferred the phylogenetic position of 43 previously unsampled Neotropical species from COI, H3, WNT, 18S, 16S, and 12S. Based on our phylogenies we found that Neotropical psocids are generally not closely related to morphologically similar taxa in the Holarctic region. Consequently, the monophyletic status for the major groups within Psocidae (subfamilies and tribes) is recovered only when Holarctic groups are sampled (7–10 of 11 higher-level groups are monophyletic) but violated when Neotropical species are included in the dataset (1 of 11 higher-level groups are monophyletic). Leveraging the largest phylogeny of the Psocidae, our study pinpoints the downfalls of simply extending taxonomic knowledge from lineages of a certain area to inform diversity and evolution of lineages in other regions.

**Highlights:**

- Tropical psocids comprise >60% of the extant family richness
- Previous phylogenies have undersampled Tropical psocids
- Holarctic and Neotropical species are classified under the same morphological groups
- Holarctic and Neotropical generally correspond to evolutionarily distinct lineages
- Phylogenies based on Holarctic psocids poorly inform evolution in the Neotropics

## 1 INTRODUCTION

### 1.1 Background

With more than 1,000 species classified across 80 different genera, Psocidae is the largest extant family of free-living lice (Psocodea: ‘Psocoptera’; Mockford, 1993; Johnson et al. 2020; Lienhard and Smithers, 2002). Although many temperate species are currently described (Lienhard and Smithers, 2002; Johnson et al. 2020), more than 60% of the family diversity is restricted the tropics (Text S1; Table S1). Nevertheless, diversity in the tropics is likely being greatly underestimated (e.g. Aldrete and Román-P, 2015; Román-Palacios et al. 2016; Oliveira et al. 2017). Species within the Psocidae are classified under three subfamilies and ten tribes (Johnson et al. 2020; Lienhard and Smithers, 2002; Yoshizawa and Johnson, 2008). Kaindipsocinae accounts for 36 species (Yoshizawa, 1998; Yoshizawa et al. 2011; Johnson et al. 2020), Amphigerontiinae includes 235 species classified into three tribes (Amphigerontini, Blastini, and Stylatopsocini; Yoshizawa 2010), finally, Psocinae, the largest clade within the Psocidae, includes nearly ~1,000 species classified under seven tribes (‘Ptyctini’, Psocini, Atrichadenotecnini, Sigmatoneurini, Metylophorini, Thyrsophorini, and Cycetini; Yoshizawa and Johnson, 2008). Although many groups within the Psocidae were included based on morphology (Yoshizawa 2002, 2005), the recent use of molecular data to study the Psocidae systematics has provided new insights on the natural groups within the family (e.g. Yoshizawa and Johnson, 2008).

A handful of molecular studies have examined the phylogenetic relationships among several higher-level groups (subfamilies and tribes) within the Psocidae. For instance, Johnson and Mockford (2003) recovered the family-level monophyly and concluded the paraphyletic status of the Psocinae based on four gene regions (18S, 12S, 16S, and COI) sequenced from four Psocidae species (three Psocinae and a single Amphigerontiinae). More recently, Yoshizawa and Johnson (2008) presented the most comprehensive species-level phylogeny for the Psocidae published to date based on six gene regions (18S, 16S, 12S, COI, H3, and ND5) and 45 Psocidae species. Relative to the morphology-based classical taxonomy (Lienhard and Smithers, 2002), Yoshizawa and Johnson (2008) erected a new tribe (Kaindipsocini), synonymized the Oriental Cerastipsocini (*Sigmatoneura* and *Podopterocus*) within Sigmatoneurini, and transferred the remaining Neotropical Cerastipsocini into Thyrsophorini. Yoshizawa and Johnson (2007) also recovered the monophyly of Psocidae and the paraphyly of both Amphigerontiinae (due to the position of Kaindipsocini; but see below) and ‘Ptyctini’. In a follow up study by Yoshizawa et al. (2011), the taxonomic sampling for Kaindipsocini in Yoshizawa and Johnson (2008) was expanded to six new species. Yoshizawa et al. (2011) also re-defined the taxonomic limits within Amphigerontiinae by limiting this subfamily to only two tribes (Amphigerontini and Blastini) and erecting a new subfamily (Kaindipsocinae, previous Kaindipsocini).

The systematics and evolution of Tropical psocids has been historically understood from studies mainly sampling on Holarctic lineages. For instance, tropical lineages represented 25% of the species sampled in Johnson and Mockford (2003; 1 of 4 taxa), ~17% in Yoshizawa and Johnson (2008; eight of 45), and ~16% in Yoshizawa et al. (2011; eight of 51). This discrepancy regarding the geographical bias of lineages sampled in molecular phylogenies (e.g. Holarctic groups) questions the practical utility of previous phylogenetic hypotheses in informing the evolution and diversity outside of the main target region (e.g. the Neotropics).

### 1.2 Objectives

When combined with classical morphological taxonomy, phylogenies generate predictions about the evolutionary position of lineages that are not sampled in trees (Hennig, 1999; Felsenstein, 2004). Here, we use the Psocidae to test whether phylogenetic trees strongly based on Holarctic species can predict the phylogenetic position of Neotropical taxa. In this study, we sequenced three gene regions for 43 Neotropical taxa that were not sampled in previous molecular phylogenies. We then inferred the phylogenetic relationships among psocid species using molecular dataset including (i) species previously sampled in studies of the Psocidae phylogeny, and (ii) Neotropical species that were generated in this study.

We expected species in the same genera, tribes, and subfamilies (originally classified based on morphology) to be closely related in the Psocidae phylogeny regardless of their geographical origins. We suggest that, because taxonomy has been largely based on morphology, and morphological convergence has shown to be widespread in the Psocidae, phylogenetic hypotheses with sampling biased towards Holarctic lineages cannot inform the phylogenetic position and diversity of Tropical lineages.

## 2 MATERIAL AND METHODS

### 2.1 Overview of molecular databases

We constructed two molecular data sets to study the Psocidae systematics. First, we generated molecular for 43 Neotropical psocid species that have not been sampled in previous phylogenetic studies. Second, we combined the newly obtained sequences with publicly available sequences that previous studies have used to infer the phylogenetic relationships within the Psocidae.

### 2.2 Field work, DNA extraction, amplification, and sequencing

We obtained molecular data for 43 species never included before in phylogenetic studies, collected from five localities in Colombia where extensive psocopteran collections have been conducted over the last decade: (1) Dagua: El Queremal, Vereda La Elsa (03°33’55.8”N; 76°45’30.0”W; (2) Cali: Los Yes, Quebrada Honda (3°26’01.8”N; 76°38’40.3”W), (3) Cali: La Buitrera (3°32’14.1”N; 76°45’19.0”W; (4) Dagua: Km 23, Via a Buenaventura, El Canasto (3°33’13.5”N; 76°36’34.6”W), y (5) Dagua: Km 18, Via a Zingara (3°32’0.1”N; 76°36’35.1”W). All collected individuals were dry-stored in vials at −4°C. Morphological identification was conducted using published taxonomic keys (e.g. Smithers, 1990) and recently published diagnoses (e.g. García-Aldrete and Román-P., 2015; Román-P. et al. 2014; Yoshizawa, 1998). All voucher specimens used in this study are deposited in the Psocopteran collection of the Universidad del Valle, Colombia (Grupo de Investigaciones Entomológicas).

We followed Birungi and Munstermann (2002) for the DNA extraction protocol (with an incubation period of one hour in potassium acetate; Rosero et al. 2010) and Ruíz et al. (2010) for reagents concentrations used in PCR. We amplified three gene regions corresponding to one mitochondrial and two nuclear genes (Table S2). PCR thermal cycle protocols used to amplify each gene region are summarized in Table S3. Sequencing was conducted in Macrogen Inc and Geneious 7.1.3 (Kearse et al. 2012) was used to assemble the final sequences.

### 2.3 Retrieval of published sequences

In addition to the newly generated sequences, we obtained molecular data on COI, 18S, and H3 genes from GenBank (Benson et al. 2012) and BOLD Systems (Ratnasingham and Hebert, 2007). We also used public databases to expand the molecular sampling in our study by including 12S, 16S, and Wingless genes. These last three genes have been extensively sampled in previous studies of the Psocidae phylogeny (e.g. Yoshizawa, 2001, 2004; Bess and Yoshizawa, 2007; Yoshizawa and Johnson, 2008; Bess et al. 2014). Additionally, the gene sequences of two outgroup free living lice species in the Hemipsocidae (*Hemipsocus chloroticus*) and Psilopsocidae (*Psilopsocus malayanus*) were sampled from publicly available databases.

### 2.4 Assembly and curation of molecular datasets

We constructed two molecular datasets for the Psocidae by assembling DNA alignments from the (i) sequences obtained through public databases, and (ii) the combination of both newly generated and published sequences. The assembly and curation of each of these two datasets was conducted following protocols based on SuperCRUNCH version 1.0 (Portik and Wiens, 2020).

We first combined all the dataset-specific sequences in a fasta file with sequence names according to SuperCRUNCH. We removed duplicated sequences (script Remove_Duplicate_Accessions) and subspecies or ambiguously identified taxa (e.g. sp., aff.; Fasta_Get_Taxa script). Next, loci-specific fasta files were generated (Parse_Loci script) based on the following alternative versions of each gene: COI (COI, COX, COX, and cytochrome), H3 (H3 and Histone 3), wingless (wingless and Wnt), 18S, 12S, and 16S. For each locus, we selected the longest sequence per species (Filter_Seqs_y_Species). We then used CD-HIT version 4.6.8 within the EST package (Li and Godzik, 2006) and BLAST (megablast; Madden, 2013) to test for the sequence orthology within each of the species-level fasta files. For each locus, we kept the largest cluster of orthologous sequences (Cluster_Blast_Extract.py script) and adjusted the direction of all sequences before performing sequence alignment under MAFFT v. 7 (Adjust_Direction script in SuperCRUNCH; Katoh and Standley, 2013). Our phylogenetic analyses are based on these orthologous clusters within each of the two datasets.

### 2.5 Sequence alignment

We used SuperCRUNCH to obtain six orthologous gene clusters from each dataset (published sequences and combined sequences). Each of these gene clusters was then aligned using the profile alignment routine implemented in MAFFT v. 7 (Katoh and Standley, 2013). For each sequence alignment in MAFFT we (i) allowed sequence direction to be adjusted, (ii) aligned length to remain the same as in the existing alignment (--add parameter), and (iii) conducted a local alignment under the L-INS-1 strategy. The remaining parameters were set to default. We selected the following set of published alignments to guide the alignment of our sequences. For COI and 12S genes, we used the alignments in Chesters (2017). We used the H3 sequence alignment from Gamboa et al. (2019). For Wingless, we followed the alignment from Phillips et al. (2017). Finally, we aligned both 16S and 18S genes by following the secondary structure indicated in Viale et al. (2015) and Kjer (2004), respectively. The sequence alignment in Kjer (2004) for 18S was transformed from RNA to DNA using Seqotron. Finally, we removed sequences that did not overlap with the regions sampled in the existing alignments (Kjer, 2004; Viale et al. 2015; Chesters, 2017; Phillips et al. 2017; Gamboa et al. 2019). We obtained a single concatenated alignment for each dataset (File S1, published sequences; File S2, combined sequences). These concatenated alignments based on profile alignments of individual loci were then used in the phylogenetic inference steps.

### 2.6 Partitioning strategies of the supermatrixes

We obtained one concatenated dataset for published sequences and another for the combined sequences. Given that the analyzed partitioning strategy of the dataset can affect the resulting phylogenetic relationships among species within each dataset, we conducted independent analyses based on alternative partitioning strategies. A partition strategy corresponds to the sequence blocks in an alignment that are set prior to a statistical analysis of the optimal partitioning (e.g. using PartitionFinder; Lanfear et al. 2017). We therefore used two partitioning strategies for each dataset: (i) gene-based partitioning, and (ii) codon/gene-based partitioning to examine optimal partitioning schemes. A partition scheme results from statistically evaluating partition strategies (results of PartitionFinder).

We run PartitionFinder twice in each dataset using two partitioning strategies that resulted in the same number of partitioning schemes per dataset. First, we used gene-based partitions within each concatenated alignment. Alternatively, we used a combination of gene-based (for the non-protein-coding genes 12S, 16S, and 18S) and codon-based (for protein-coding genes COI, H3, and wingless) partitioning for each dataset. PartitionFinder output files are provide in File S3.

### 2.7 Phylogenetic analyses

We obtained two different partitioning schemes for each of the two molecular datasets. We followed Baca et al. (2017) to compare the fit of these partitioning schemes. Phylogenetic inference was performed under Maximum Likelihood in RAxML-HPC BlackBox 8.2.10 (Stamatakis, 2014) and Bayesian Inference in MrBayes (Ronquist et al., 2012). We run all phylogenetic analyses in CIPRES Science Gateway V. 3.3 (Miller et al. 2010). Under RAxML, we set a total of 1,000 bootstrap replicates, used a GTRGAMMA model for each partition, and set the remaining parameters to default. Under MrBayes, we performed two simultaneous runs for each combination of dataset and partitioning scheme consisting of eight MCMC chains (one cold and seven heated) chains running for 30 million generations. Trees were sampled every 1,000 generations. We assessed convergence of parameters by investigating the Effective Sample Size (ESS) of all parameters in Tracer 1.7 (Rambaut et al, 2018). A value of ESS > 200 was indicative of convergence. We discarded 10% of posterior trees as burn-in and inferred the 50% majority rule consensus tree based on the remaining samples. Finally, we compared the performance of partitioning schemes based on likelihood estimates from RAxML runs. The best partitioning scheme for each dataset was selected based on the highest likelihood score under maximum likelihood in RAxML.

## 3 RESULTS

We sequenced three gene regions from 43 Neotropical psocopteran species in the Psocinae and Amphigerontiinae (Table 1). Four of these samples were not morphologically similar to any of the currently described tribes and subfamilies in the Psocidae. To our knowledge, all species that were sequenced in this study are exclusively restricted to the Neotropics.

**Table 1.**
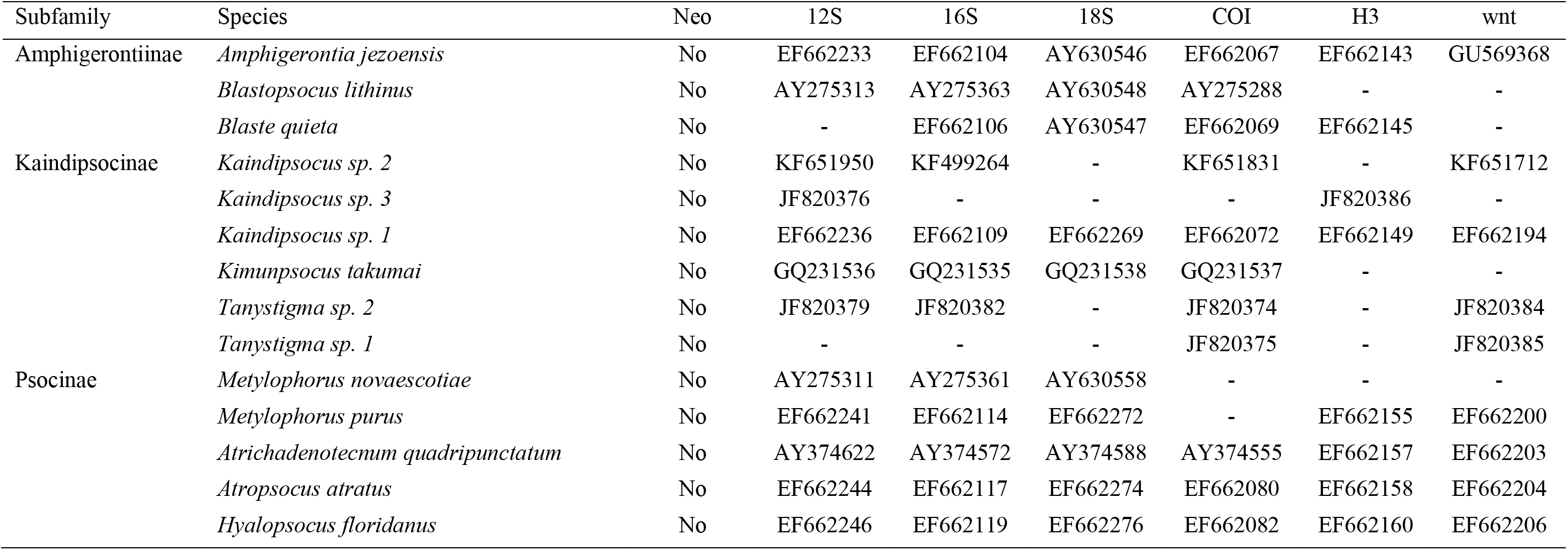
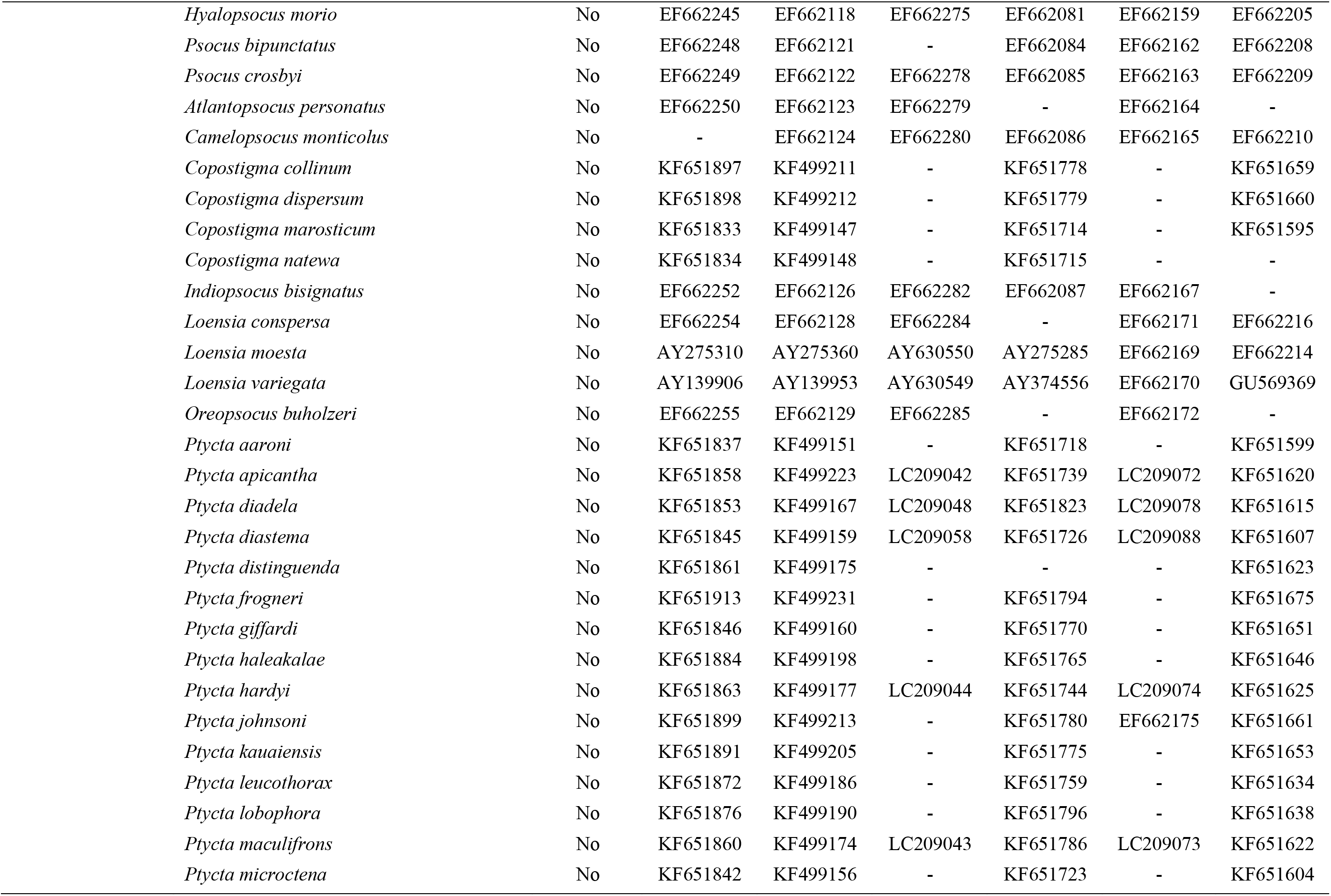
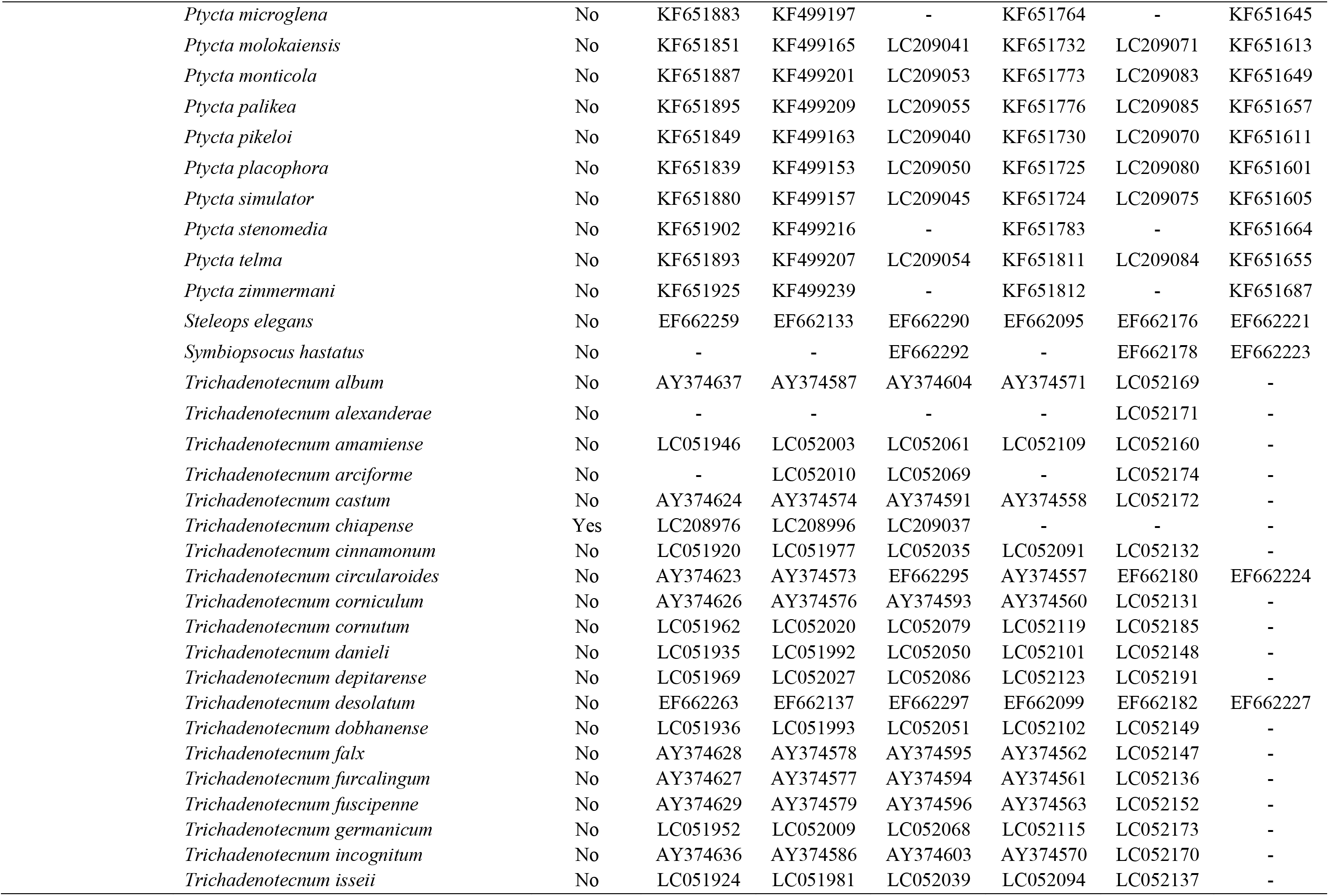
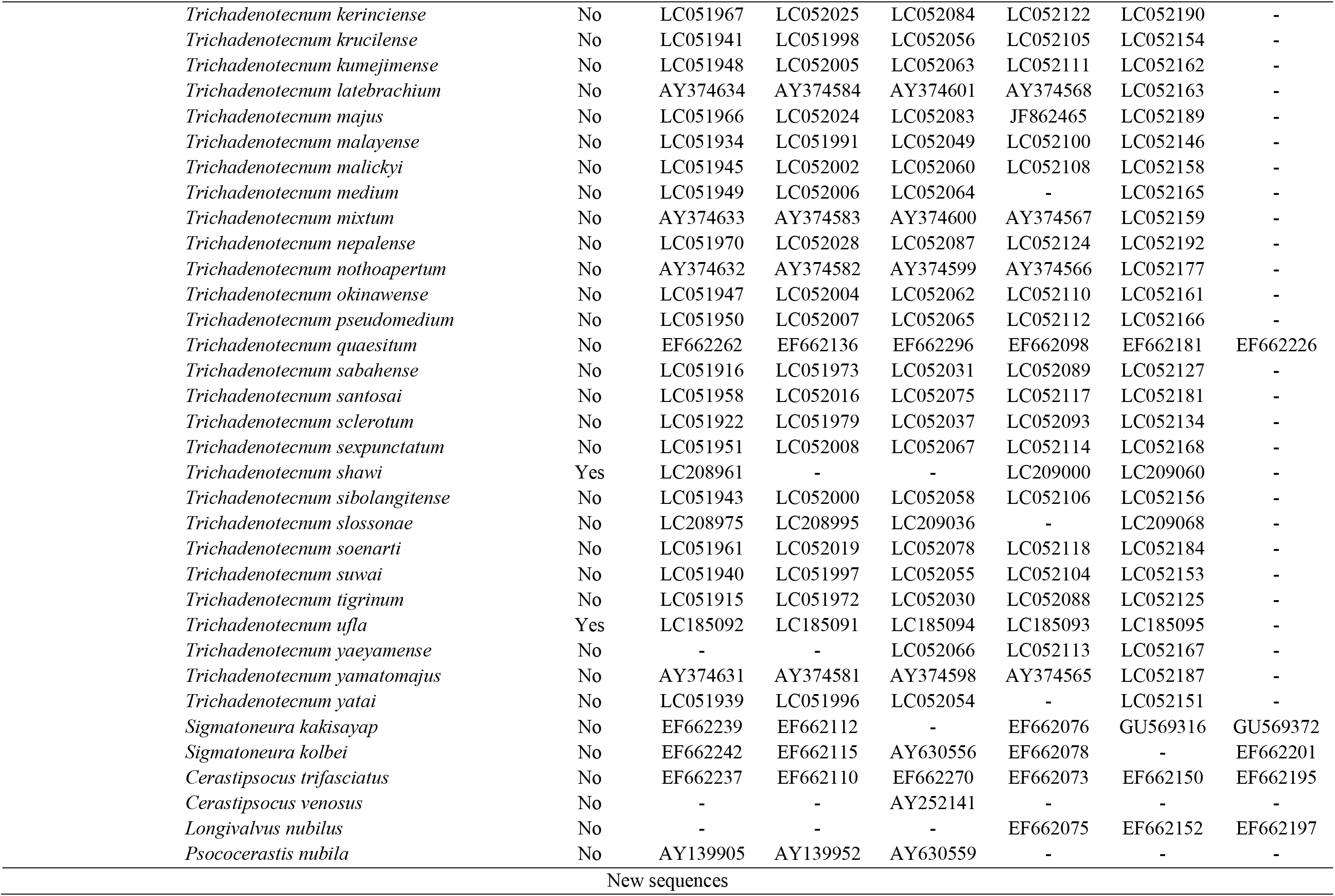
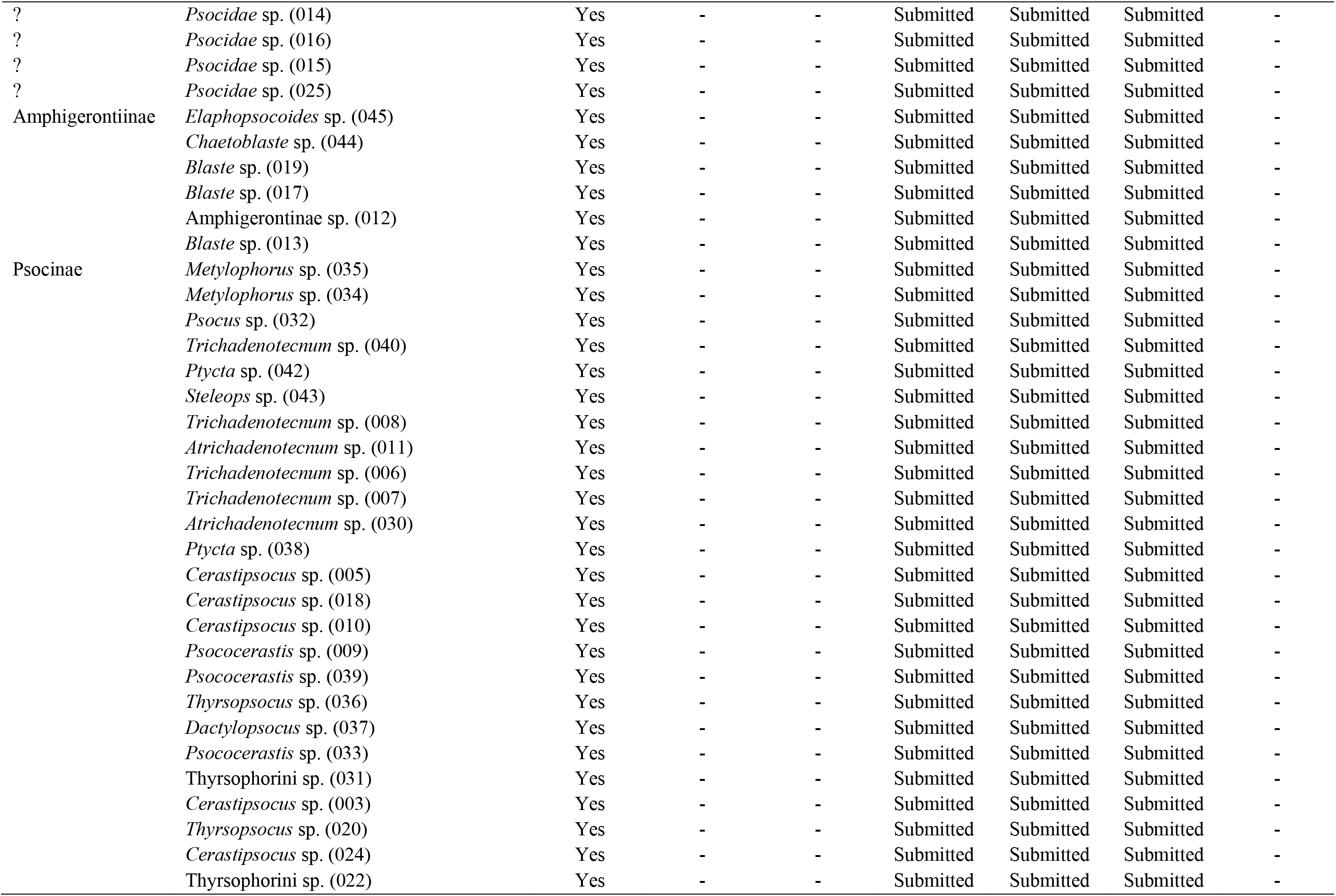
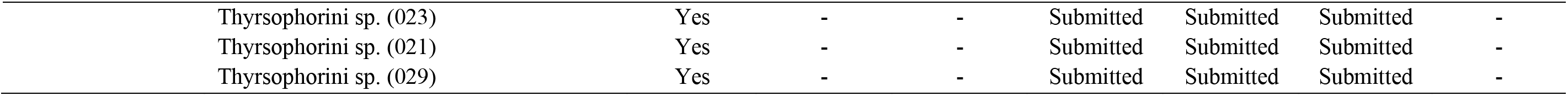
Molecular sampling and geographical distribution of the specimens examined in this study. We summarize the taxonomic information for each species, along with information on whether its distribution includes Neotropical areas or not (column=Neo). Samples obtained in this study are indicated on the bottom of the table. Each of these samples is labeled with a code corresponding to the individual number deposited in the Psocopteran collection of the Universidad del Valle, Colombia (Grupo de Investigaciones Entomológicas).

Our phylogenetic analyses were based on 38 of the total 43 Neotropical species sequenced in this study – five species were excluded in different stages of the dataset construction under SuperCRUNCH. Since our main interest was on testing if the phylogenetic position of Neotropical species could be predicted from a strongly Holarctic-biased phylogeny, we generated two datasets. First, we retrieved from public databases all available sequences for the Psocidae. Second, we combined our newly generated sequences with published sequences. In total, our published-sequences dataset included 109 Psocidae species from 25 genera, eight tribes, and three subfamilies. The combined dataset included 147 Psocidae species from 30 genera, eight tribes, and three subfamilies. The phylogenetic relationships among the species in each of these datasets was analyzed under two different partition schemes. Our main results for both Maximum Likelihood analyses and Bayesian Inference trees are based on the partitioning scheme resulting in the higher likelihood (under Maximum Likelihood). Specifically, a codon-based partitioning strategy was selected as the best-fitting approach (i.e. the model with the highest likelihood under Maximum Likelihood) for both the combined (Codon=−48142.629, Genes=−49088.281) and public datasets (Codon=−42862.351, Genes=−43553.692). Nevertheless, all trees recovered similar phylogenetic relationships among lineages.

Phylogenetic analyses based on the published-only dataset inferred the family-level monophyly (bootstrap=100%; Fig. 1). At the subfamily level, our analyses recovered the monophyly for Amphigerontiinae (bootstrap=100%) and indicated paraphyly for Kaindipsocinae and Psocinae. Specifically, *Kimunpsocus takumai* (Kaindipsocinae) and multiple Psocinae (*Oreopsocus buholzeri, Loensia conspersa, Camelopsocus monticolus, Loensia variegata*, and *Loensia moesta*) were found to cluster outside the core clades of each these two groups. The species causing the paraphyly of Psocinae and Kaindipsocinae were consistently recovered as being closely related to Amphigerontiinae. At the level of tribes, our analyses recovered (but sometimes weakly supported) the monophyly of Blastini (bootstrap=65%), Metylophorini (bootstrap=53%), and Sigmatoneurini (bootstrap=100%). Our analyses did not test the Amphigerontiini monophyly (we only sampled *Amphigerontia jezoensis*). Pyctini was recovered as a paraphyletic group, with several Pyctini being found closely related to species in almost every other tribe in the Psocinae. We note that although our analyses recover the monophyly of *Trichadenotecnum* and *Ptycta* + *Copostigma* (bootstrap=83% and 100%, respectively), our results do not support a clade including these three genera: *Trichadenotecnum*, *Copostigma*, and *Ptycta* (bootstrap=1%). *Atrichadenotecnum* was found to cluster with *Trichadenotecnum*, *Copostigma*, and *Ptycta,* but this clade was not supported (bootstrap=4%). We recovered the paraphyly of Psocini, with several Psocini being found closely related to species in a clade formed by Sigmatoneurini + Thyrsophorini + Metylophoriny (bootstrap=14%). Finally, we inferred the paraphyly of Thyrsophorini caused by *Longivalvus nubilus* closely related to Sigmatoneurini. We recovered a core clade of Thyrsophorini comprising all *Cerastipsocus* and *Psococerastis* species in our dataset (bootstrap=63%).

**Figure 1.**
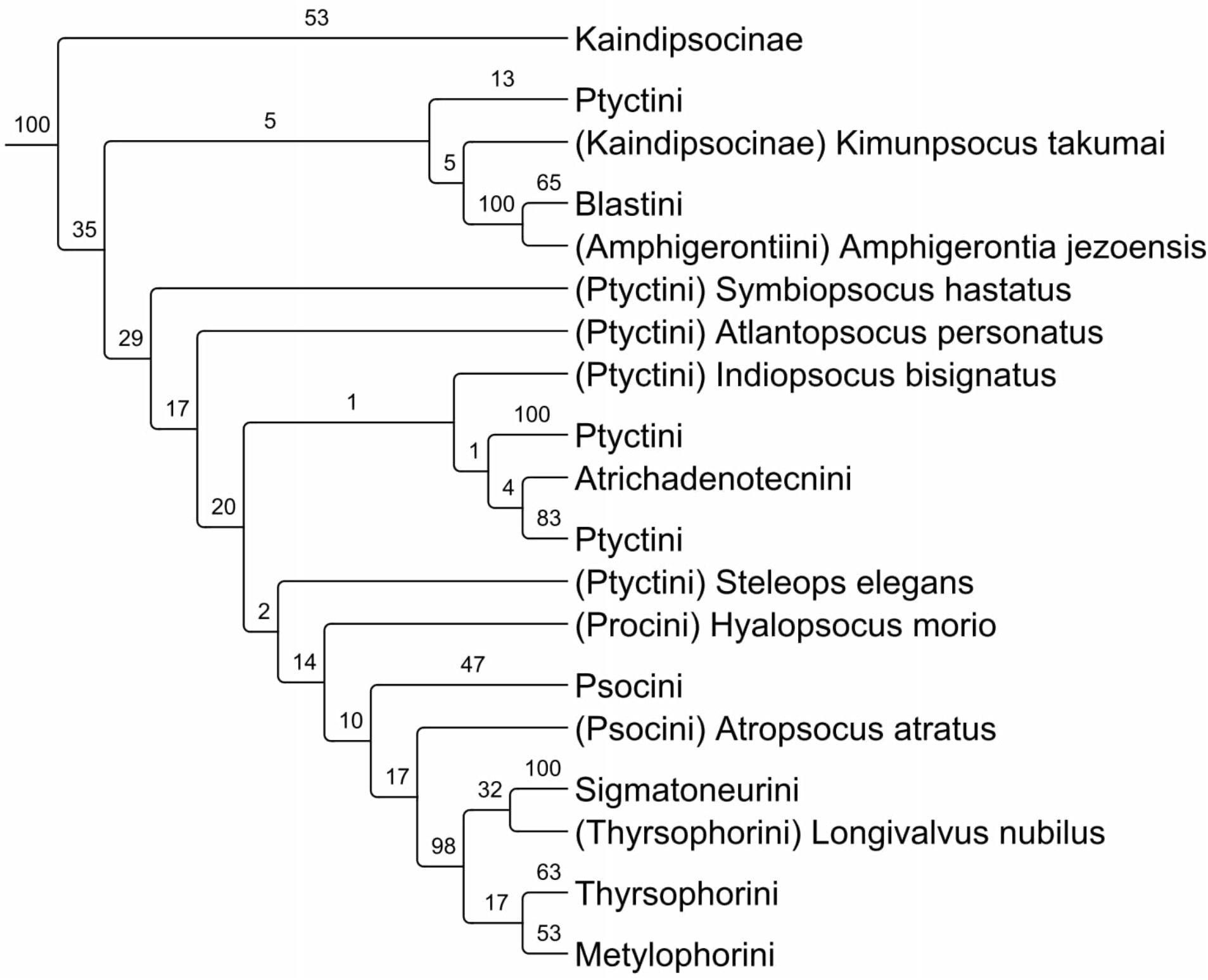
Phylogenetic relationships among the Psocidae based on the published-only dataset and reconstructed using Maximum Likelihood using a codon-based partitioning scheme. We summarize the higher-level taxonomy for the species in the tree. The full species-level phylogeny is presented in Supplementary Information S4, but additional results under alternative partitioning schemes of the alignment are included in the Supplementary Information S5. Results based on Bayesian analyses for the same dataset are similarly included in the Supplementary Information S8–S9.

We then examined the phylogenetic relationships within Psocidae based on a second dataset expanding the species-level sampling of the published-only dataset by including 38 Neotropical species. Based on the combined dataset, we did not infer the monophyly for any of the three subfamilies (Fig. 2). Within Kaindipsocinae, the phylogenetic closeness of *Kimunopsocus* to several Ptyctini generated the non-monophyly of the subfamily (bootstrap=28%). Within Amphigerontini, *Elaphopsocoides* was found as sister to all the Psocidae (bootstrap=70%) and *Amphigerontia* was found in the core Amphigerontiinae (bootstrap=42%). Finally, species in Psocinae clustered with species from the other two subfamilies. At the tribal level, we only recovered the monophyly for Sigmatoneurini (bootstrap=100%). Within Blastini, Neotropical *Blaste* were closely related to *Amphigerontia* (bootstrap=39%) and *Chaetoblaste* to *Metylophorus* (bootstrap=60%). The monophyly of Amphigerontiini was also rejected due to the position of *Elaphopsocoides* as sister to the rest of Psocidae (bootstrap=64%). Within Pyctini, all the Holarctic *Trichadenotecnum* were still recovered forming a monophyletic group (bootstrap=34%). However, we recovered two Neotropical *Trichadenotecnum* in a second clade (including *Atrichadenotecnum* and *Indiopsocus*) that was sister to the remaining *Trichadenotecnum* (bootstrap=29%). Although Holarctic *Ptycta* and *Copostigma* formed a well-supported clade (bootstrap=95%), not all Neotropical *Pycta* were clustered within this group. Within Metylophorini, one of the two species sampled in our dataset clustered with Neotropical taxa from *Ptycta*, *Psococerastis*, and *Chaetoblaste* (bootstrap=31%). Our analyses inferred the monophyly of all non-Neotropical species of Psocini (bootstrap=42%) but placed a Neotropical species of Psocini as closely related to Neotropical *Trichadenotecnum* (bootstrap=93%). Finally, although most Thyrsophorini formed a single clade that also included two Neotropical *Metylophorus* (bootstrap=43%), *Longivalvus nubilus* was sister to Sigmatoneurini (bootstrap=60%) and a single *Psococerastis* closely related to a Neotropical *Ptycta* (bootstrap=100%).

**Figure 2.**
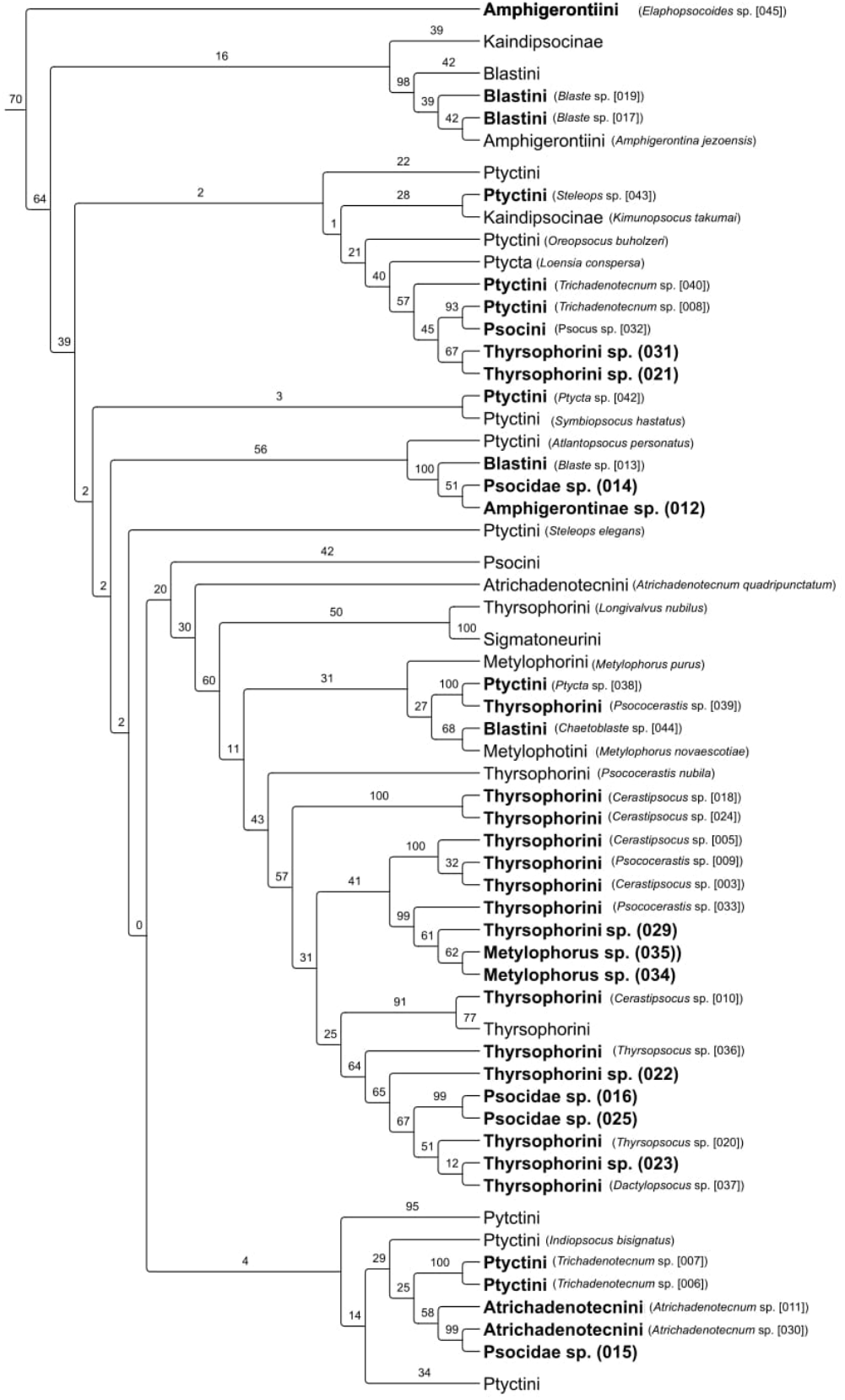
Phylogenetic relationships among the Psocidae based on the combined dataset and reconstructed using Maximum Likelihood using a codon-based partitioning scheme. We summarize the higher-level taxonomy for the species in the tree. However, we present each of the Neotropical species sequenced in this study as independent tips with tip names boldfaced and codes corresponding with those presented in Table 1. The full species-level phylogeny is presented in Supplementary Information S6, but additional results under alternative partitioning schemes of the alignment in the Supplementary Information S7. Results based on Bayesian analyses for the same dataset are shown in the Supplementary Information S10–S11.

## 4 DISCUSSION

Leveraging the most comprehensive species-level molecular dataset for the Psocidae (147 species or three times larger than previous phylogenies with 45 species), we inferred the phylogenetic relationships among all extant subfamilies, 80% tribes (8 of 10), and ~38% of genera in the family (30 of ~80). Relative to recent studies on the Psocidae phylogeny, our study increases the sampling of Neotropical taxa in the Psocidae phylogeny by a factor of ~5 (from eight species in the most recent Psocidae phylogeny (Yoshizawa and Johnson, 2008) to 38 species in our study. Nevertheless, we acknowledge that conclusions on the systematics within the family are still unreliable given the size of our phylogeny in the relation to the total family diversity (15% of ~1000 species). Our study represents an interesting case study for lineages in which (i) most morphological and molecular studies have been conducted on Holarctic taxa, and (ii) where the systematics of Tropical lineages is understood from morphological resemblance to taxa in other regions. Below, we discuss the implications of our findings on the Psocidae Tree of Life in the context of previous phylogenetic hypotheses.

### 4.1 Can heavily-Holarctic sampled phylogenies predict the phylogenetic position of Neotropical taxa?

Relative to recent phylogenies of the Psocidae (e.g. Yoshizawa and Johnson, 2008), we inferred similar phylogenetic relationships within and between taxa in Kaindipsocinae, Ptyctini, Psocini, Thyrsophorini, Sigmatoneurini, Metylophorini, Amphigerontiini, and Blastini (Figs. 1, 3). Nevertheless, the inclusion of Neotropical taxa (Fig. 2) had major changes to the relationships within and among major groups within the Psocidae (Figs. 1, 3, 4). Neotropical species in *Elaphopsocoides*, *Psocus* (032), *Blaste* (013), *Ptycta* (038), *Psococerastis* (039), *Chaetoblaste* (044), *Metylophorus* (029, 035, 034), *Trichadenotecnum* (006, 007), and *Atrichadenotecnum* (011, 030) did not cluster within their corresponding morphological groups. In general, this incongruence in the systematics of the Psocidae is likely caused by the historical undersampling of Neotropical taxa in previous phylogenetic studies (e.g. Mockford, 1993; Yoshizawa and Johnson 2008; Liu et al. 2013). The fact that the evolutionary history in the Psocidae is currently mostly understood from Holarctic lineages, neglects the description of a potentially higher diversity in the tropics.

**Figure 3.**
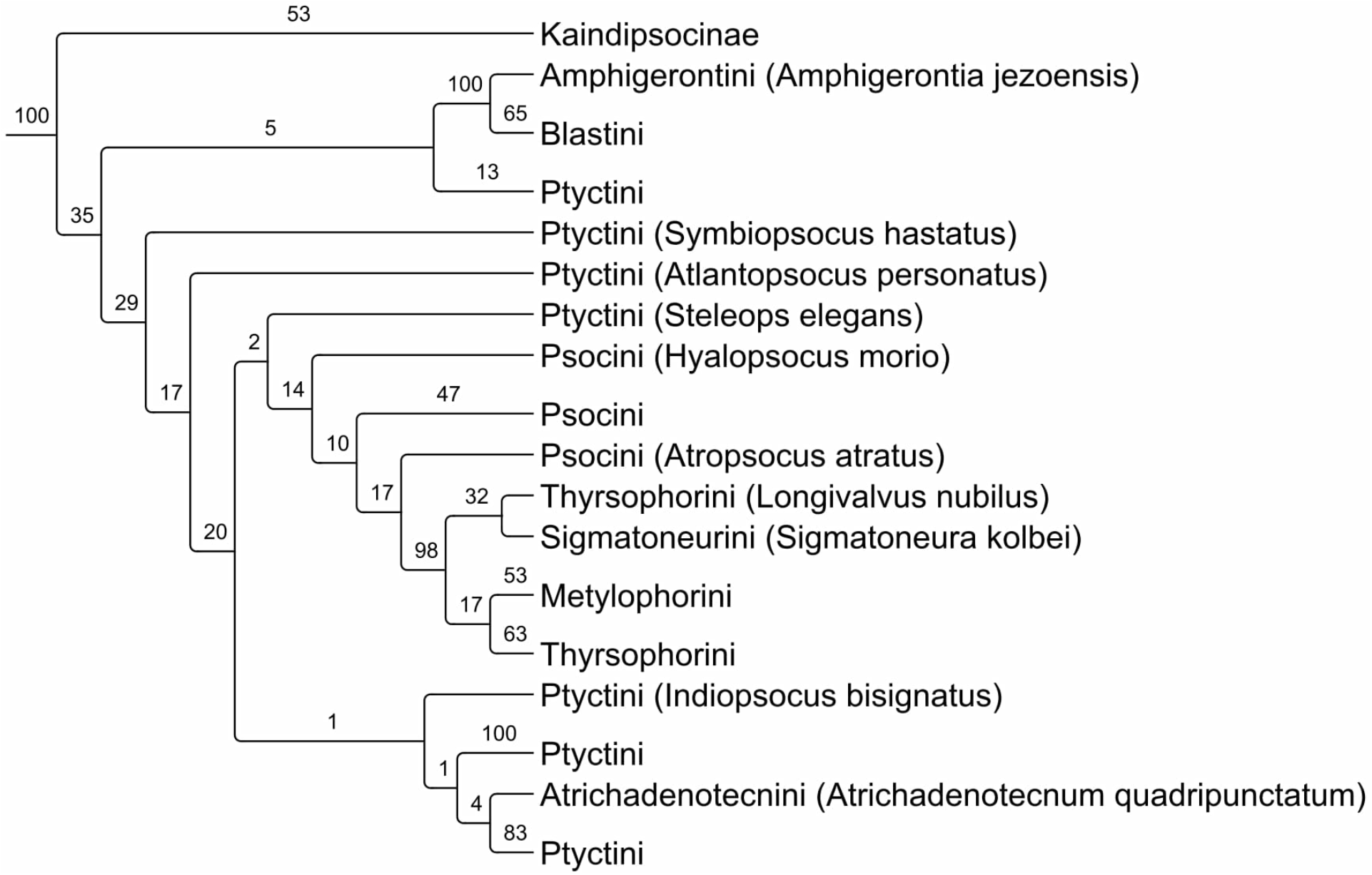
Phylogenetic relationships among the Psocidae based on the published-only dataset and reconstructed using Maximum Likelihood under a codon-based partitioning scheme. This figure is equivalent to Fig. 1 in this document but excluding species that are not shared with a previous study on the Psocidae phylogeny (Yoshizawa and Johnson 2008). The full species-level phylogeny is presented in Supplementary Information S4, but additional results under alternative partitioning schemes of the alignment are included in the Supplementary Information S5. Results based on Bayesian analyses for the same dataset are similarly included in the Supplementary Information S8–S9.

**Figure 4.**
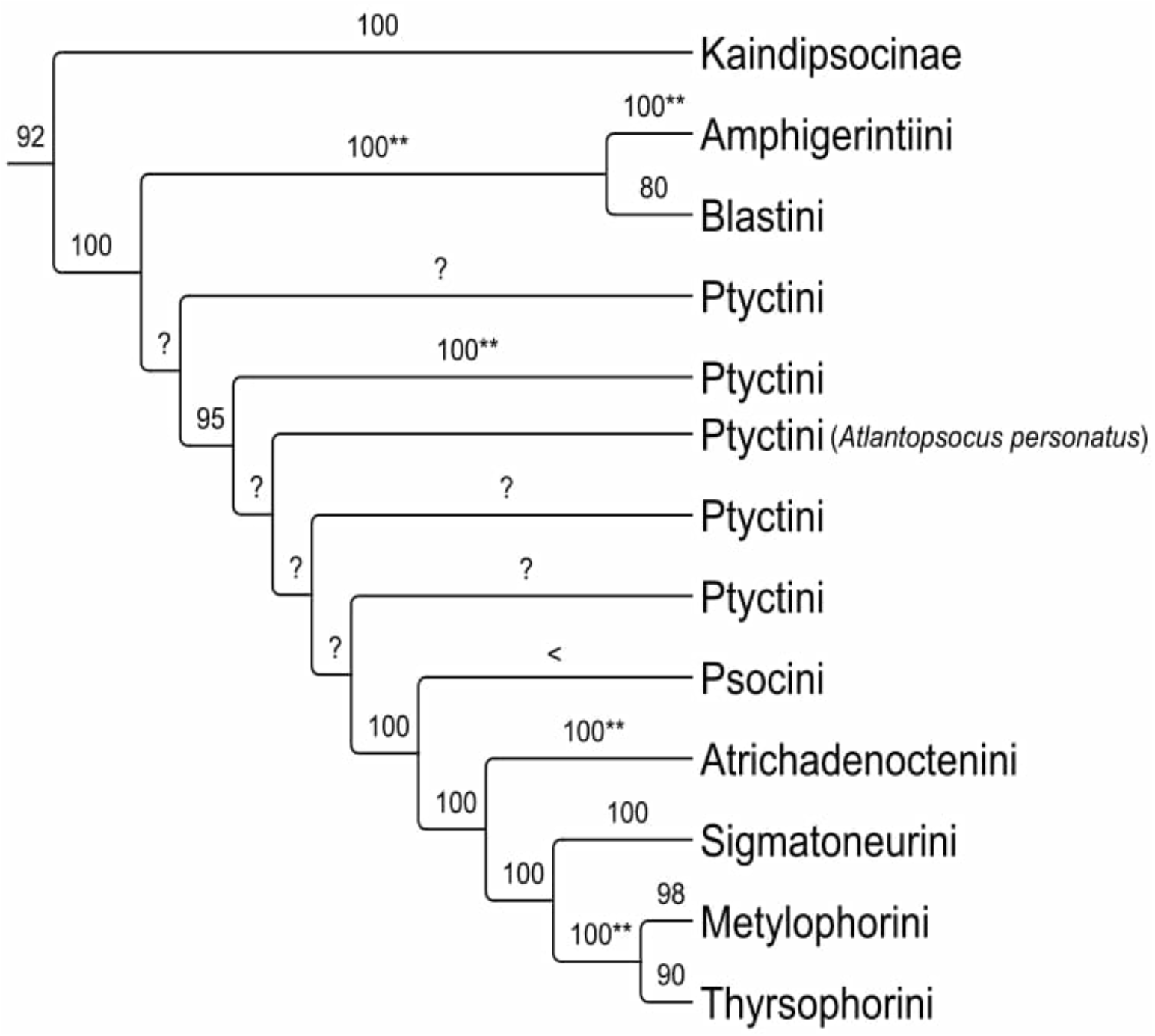
Higher-level relationships within the Psocidae modified from Yoshizawa and Johnson (2008). We present the major groups recovered in the same study. Support values based on Maximum Likelihood from the same study are indicated in the figure. We also present the following additional codes summarizing relevant aspects of their phylogenetic analyses: “**” Enforced monophyly, “?” No support values provided in the tree, “<” low support value not shown.

### 4.2. What can we learn from previous molecular phylogenies of the Psocidae Tree of Life?

To our knowledge, Yoshizawa and Johnson (2008) present the most comprehensive species-level phylogeny of the Psocidae published to date. In that study, the authors sampled 45 taxa from almost all the major groups within the Psocidae and discussed the systematic status of the same higher-level lineages. Two aspects, however, potentially obscured the importance and extent of their phylogenetic conclusions (Fig. 4). First, the monophyletic status for at least four groups (i.e. Amphigerontiini, several Ptyctini, Atrichadenoctenini, and Metylophorus + Thyrsophorini) was enforced without prior testing (but see topological tests applied for Psocini [monophyly not rejected], Amphigerontiinae [monophyly not rejected], Psocinae [monophyly rejected], Ptyctini [monophyly rejected], and Metylophorini [monophyly not rejected]). Topological constraints may result in suboptimal trees (Maddison et al. 1998; Möller et al. 2018). Second, despite a fully bifurcating phylogeny being presented in Yoshizawa and Johnson (2008), the relationships within Psocinae are very ambiguous given the lack of support values for many of the groups that are shown as being apparently resolved in the tree (Fig. 4). Future studies on the Psocidae phylogeny should also clearly and explicitly highlight their limitations.

### 4.3 Morphological convergence: Morphological vs molecular phylogenetics in the Psocidae

Our analyses indicate that morphological taxonomy largely disagrees with molecular systematics in the Psocidae. Out of the three subfamilies (Kaindipsocinae, Amphigerontiinae, Psocinae) and seven tribes (Amphigerontiini, Blastini, Psocini, Atrichadenoctenini, Sigmatoneurini, Metylophorini, and Thyrsophorini) recovered as monophyletic in previous studies mostly based on Holarctic taxa (e.g. Yoshizawa and Johnson, 2008; Yoshizawa et al. 2014), only one tribe (Sigmatoneurini) was inferred as monophyletic after the inclusion of Neotropical lineages. Because our phylogenetic analyses based (i) on published data used in previous studies (Fig. 1) and (ii) the expanded dataset including more Neotropical taxa (Figs. 2–3), did not significantly differ from published phylogenies (Figs. 3–4), we conclude that the inclusion of Neotropical taxa drove the non-monophyletic status for nine higher-level groups within Psocidae.

We found that morphological classification does not accurately reflect evolutionary closeness in the Psocidae. For instance, our analyses suggest that not all Neotropical and Holarctic *Trichadenoctenum*, a clade that has been historically highly supported by molecular and morphological data, cluster in a single clade. Similarly, *Elaphopsocoides*, an exclusively Neotropical genus (Román-P. et al. 2014), was not recovered within the remaining Amphigerontiini, a tribe that has also been inferred as monophyletic in previous studies (Yoshizawa and Johnson 2008; Yoshizawa et al. 2011). In a more striking example, Neotropical species of Methylophorini were recovered as being closely related to Thyrsophorini. However, the only Holarctic species in this tribe, *Metylophorus novaescotiae*, was found closely related to the Neotropical *Chaetoblaste* (within Amphigerontiinae: Blastini; Aldrete and Román-P. 2015). In short, morphological resemblance between Neotropical and Holarctic taxa is, in many cases, not an indicative of recent common ancestry within the Psocidae.

Finally, we note that morphological classification, which is largely based on Holarctic taxa, may have hindered a large fraction of diversity and evolutionary uniqueness of Neotropical lineages. While many Neotropical lineages correspond with morphological descriptions of Holarctic taxa, many of these Neotropical groups have an independent evolutionary origin. Multiple debates about the high frequency of morphological convergence in the Psocidae, along with other studies on problematic synapomorphies within groups, further support our conclusions. Our analyses recover many Neotropical lineages to be distantly related to their morphologically closest lineages. This pattern suggests that the diversity and evolutionary differentiation across different taxonomic levels (e.g. genera, tribes, and subfamilies) in the Tropics is potentially higher than what is currently known based on Holarctic groups.

## 5 CONCLUSIONS

We show that molecular phylogenetics and morphological taxonomy strongly based on Holarctic groups cannot inform the phylogenetic position of Neotropical taxa. In addition to calling for new classification within the Psocidae, our analyses suggest that multiple Neotropical Psocidae represent independent lineages to the ones known in the Holarctic region. Although the role geography in affecting taxonomic boundaries within clades remains largely unexplored, our results suggest that, for certain groups such as the Psocidae, morphological and phylogenetic classification based on lineages found in certain areas (e.g. Holarctic) do not reflect the evolutionary history of morphologically similar taxa in other regions (Neotropics). Future studies on the Psocidae Tree of Life should rely on a better sampling of non-Holarctic lineages to derive a comprehensive hypothesis of the systematics within the family.

## 6 ACKNOWLEDGEMENTS

We thank the Laboratorio de Estudios Ecogenéticos y Biología Molecular of the Universidad del Valle for allowing us to use their facilities and equipment to conduct molecular protocols. We are also grateful to the Department of Biology, Facultad de Ciencias Naturales y Exactas, Vicerrectoría de Investigaciones, Universidad del Valle, for their constant support. We are especially thankful for the feedback provided by Cesar Medina, Heidi S. Steiner, and Jose Sergio Hleap on different stages of this project. This study was partially funded by Fundación Banco de la República.

## SUPPLEMENTARY FILES

**Supplementary File S1.** Published-only dataset alignment.

**Supplementary File S2.** Combined dataset alignment.

**Supplementary File S3.** PartitionFinder results.

**Supplementary File S4.** Phylogeny of the Psocidae based published-only dataset, inferred under Maximum Likelihood and with gene-based partitioning.

**Supplementary File S5.** Phylogeny of the Psocidae based published-only dataset, inferred under Maximum Likelihood and with codon-based partitioning.

**Supplementary File S6.** Phylogeny of the Psocidae based combined dataset, inferred under Maximum Likelihood and with gene-based partitioning.

**Supplementary File S7.** Phylogeny of the Psocidae based combined dataset, inferred under Maximum Likelihood and with codon-based partitioning.

**Supplementary File S8.** Phylogeny of the Psocidae based published-only dataset, inferred under Bayesian Inference and with gene-based partitioning.

**Supplementary File S9.** Phylogeny of the Psocidae based published-only dataset, inferred under Bayesian Inference and with codon-based partitioning.

**Supplementary File S10.** Phylogeny of the Psocidae based combined dataset, inferred under Bayesian Inference and with gene-based partitioning.

**Supplementary File S11.** Phylogeny of the Psocidae based combined dataset, inferred under Bayesian Inference and with codon-based partitioning.

**Appendix S1**. Distribution of Psocids, primers, PCR conditions, bayesian phylogenies.

## Notes

### Competing Interest Statement

The authors have declared no competing interest.

